# Decoding reward–curiosity conflict in decision-making from irrational behaviors

**DOI:** 10.1101/2022.04.24.489304

**Authors:** Yuki Konaka, Honda Naoki

## Abstract

Humans and animals are not always rational. They not only rationally exploit rewards but also explore an environment, even if reward is less expected, owing to their curiosity. However, the mechanism of such curiosity-driven irrational behavior is largely unknown. Here, we developed a novel decision-making model for a two-choice task based on the free energy principle, which is a theory integrating recognition and action selection. The model successfully described irrational behaviors depending on the curiosity level. We then proposed a machine learning method to decode temporal curiosity from behavioral data, which enables us to quantitatively compare estimated curiosity and neural activities. By applying it to rat behavioral data, we found that the irrational choices sticking to one option was reflected to the negative curiosity level. Our decoding approach can be a fundamental tool for identifying the neural basis for reward-curiosity conflicts. Specifically, it could be effective in diagnosing mental disorders.

## Introduction

Animals and humans perceive the external world through their sensory systems and make decisions accordingly^1,2^. Generally, they cannot make optimal decisions because of the uncertainty of the environment as well as the limited computational capacity of the brain and time constraints in decision-making^3^. In fact, they perform irrational actions. For example, people play lotteries and gamble despite low reward expectations. In this case, they face a dilemma between low expected reward and curiosity regarding whether a reward will be acquired. Thus, understanding how animals control the balance between reward and curiosity is important for the elucidation of the whole decision-making process. However, no method to quantify the reward– curiosity balance has yet been established. In this study, we developed a machine learning method to decode the time series of the reward–curiosity balance from animal behavioral data.

Some irrational behaviors emerge because of the strength of curiosity^4,5^. For example, conservative individuals avoid uncertainty and prefer to select an action that leads to predictable outcomes. Conversely, inquisitive individuals strongly desire to know the environment rather than rewards and prefer to select an action that leads to unpredictable outcomes. Rational individuals fall midway between these two extremes; in an ambiguous environment, they select an action to efficiently understand the environment, and if the environment becomes clear, they select an action to efficiently exploit the rewards. Thus, curiosity has a significant impact on behavioral patterns, and it is naturally thought that animals control the balance between reward and curiosity in a context-dependent manner.

Adaptive learning behaviors have been modeled primarily by reinforcement learning (RL), which is a theory for describing reward-seeking behaviors^6^. However, animals not only exploit rewards but also explore the environment, even without rewards, to minimize the uncertainty of the environment owing to their curiosity. Recently, the free energy principle (FEP) was proposed by Karl Friston under the Bayesian brain hypothesis, in which the brain optimally recognizes the outside world according to Bayesian estimation^7–9^. FEP addresses not only the recognition of the external world but also action selection, which minimizes the uncertainty of the recognition of the external world. Furthermore, FEP was integrated with RL, and then action selection was formulated by maximizing both reward and curiosity. Note that curiosity can be regarded as information gain, that is, the extent to which we expect our recognition to be updated by new observation through the action^10,11^. However, FEP assumes that the weighting of rewards and curiosity is always constant and thus cannot treat actual animal behaviors in which the weights of rewards and curiosity are expected to change over time. Hence, conventional theories such as RL and FEP are limited in describing the conflict between reward and curiosity.

The identification of the temporal variability of curiosity is important for elucidating the neural mechanisms of the reward and curiosity conflicts in decision-making. Many neuroscience studies have extensively examined the neural mechanisms of decision-making in the context of RL^12–16^. RL has worked as a guiding principle in the quantitative understanding of reward-seeking behavior. However, RL does not focus on curiosity-driven behavior; therefore, it is difficult to quantitatively understand the reward–curiosity conflict. To overcome this difficulty, a method to decode the temporal variability of curiosity from behavioral data is essential. Such a method would enable us to analyze neural correlates with the temporal variability of curiosity and, consequently, clarify how the brain controls the balance of reward and curiosity in a context-dependent manner.

In this study, we extended FEP by incorporating a meta-parameter that controls the conflict dynamics between reward and curiosity, called the reward–curiosity decision-making (ReCU) model. The ReCU model is able to exhibit various behavioral patterns, such as greedy behavior toward reward, information-seeking behaviors with high curiosity, and conservative behaviors avoiding uncertainty. Moreover, we developed a machine learning method, called the inverse FEP (iFEP) method, to estimate the internal variables of decision-making information processing. By applying the iFEP method to behavioral time series in a two-choice task, we successfully estimated the internal variables, such as variations in curiosity, recognition of reward availability, and its confidence.

## Results

### ReCU model: decision-making with reward–curiosity dilemma in a two-choice task

Animals perceive the environment by inferring causes, such as reward availability from observation, and then make decisions based on their own inferences (**Fig. 1A**). In this study, we developed an ReCU model of a decision-making agent facing a dilemma between reward and curiosity in the case of a two-choice task, in which the agent selects either of two choices associated with the same rewards but with different reward probabilities. If the agent aims to maximize cumulative rewards, the agent must select an option with a higher reward probability. However, in animal behavioral experiments, even after they learned which option was associated with a reward, they did not exclusively always select the best choice, but also often selected the option with a smaller reward probability, which seems unreasonable. Consequently, we hypothesized the following: Animals assume that the reward probability for each option might fluctuate over time so that the continuous selection of one option decreases the confidence of the reward probability estimation for the other option. Thus, they become curious about the ambiguous option even with a smaller reward probability, and then selecting the ambiguous option is reasonable to increase the confidence of the estimation for both options. Therefore, we considered that the agent should make a decision driven by reward and curiosity in a situation-dependent manner.

**Fig. 1:**
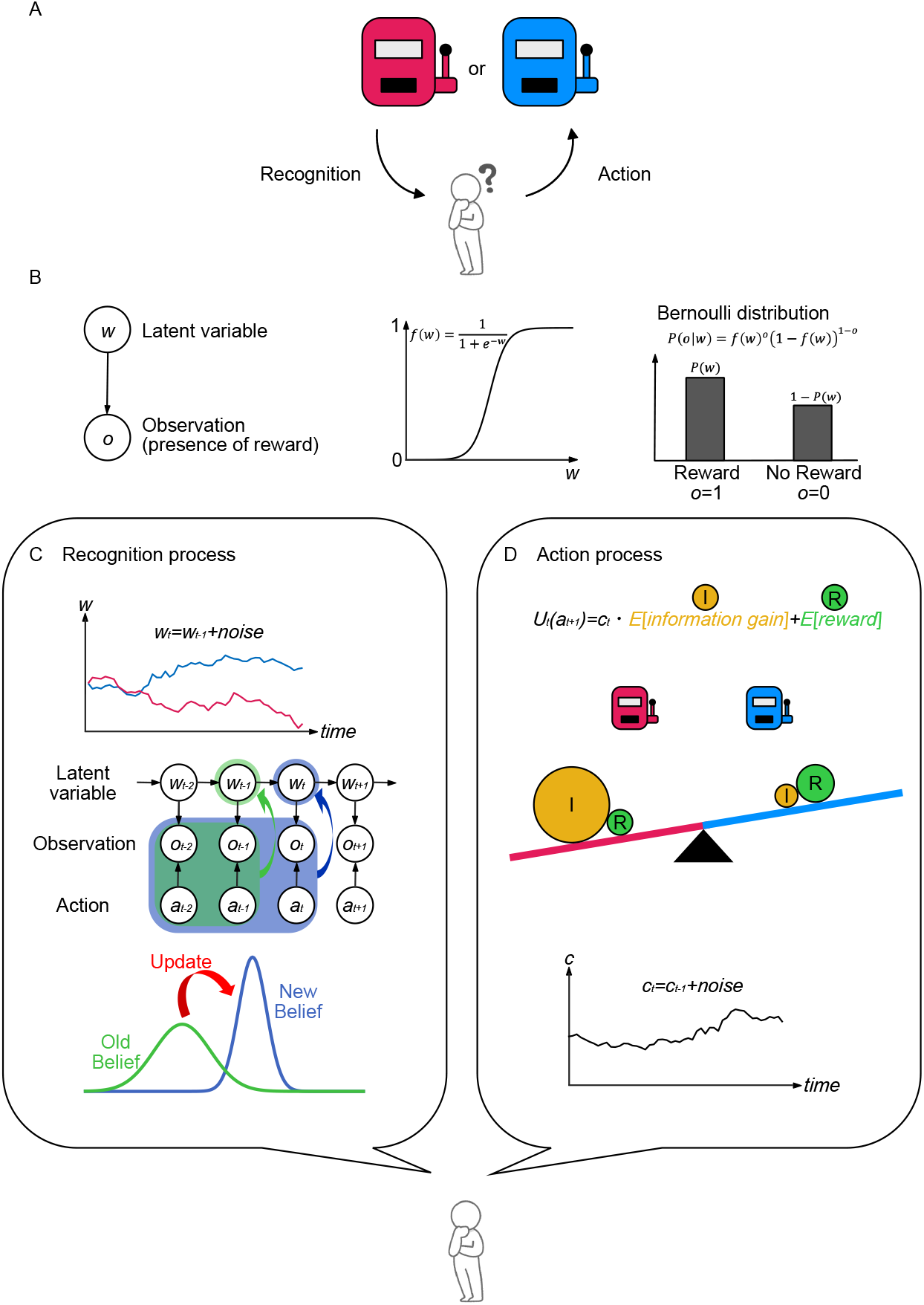
Decision-making model for the two-choice task with reward–curiosity dilemma. **(A)** Decision-making in the two-choice task. Reward is provided at different probabilities for each option. The agent does not know those probabilities. Through repeated trial and error, the agent recognizes the world by inferring the latent reward probability of each option, and decides to choose the next action, i.e., option, based on its own inference. **(B)** Generative model of reward. The agent assumes that reward is generated by the latent variable *w* (left panel), and the reward probability is sigmoidally controlled by w (middle panel). The presence of the reward obeys a Bernoulli distribution (right panel). **(C)** Recognition process by agent. The agent assumes that the reward probabilities change over time owing to fluctuation of the latent variable *w*_*t*_ (upper). The probabilistic dependence among the latent variable *w*, action *a*, and the presence of reward *o* is graphically described (middle). The agent recognizes *w* as a probability distribution, whose precision, i.e., inverse of variance, represents confidence, and sequentially updates its recognized distribution from previous rewards and actions in a Bayesian manner (lower). **(D)** Action selection process by agent. The agent evaluates the utility of each action using the weighted sum of the expected reward and information gain (upper). The agent compares the utilities for both actions and prefers the option with the larger utility (middle). The expected information gain can be calculated as the difference between the currently recognized distribution and the predicted recognized distribution updated by the future observation (lower).

In this two-choice task, the reward is given as all-or-none, but its intensity depends on the agent’s own feeling, as

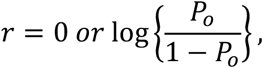

where *P*_*o*_ is a parameter that controls the reward intensity felt by the agent. According to the Friston formulation, *P*_*o*_ represents the desired level of how much the agent wants the reward^10,11^. To model decision-making driven by reward and curiosity, we divided information processing in the brain into two processes. First, the agent recognizes the world, that is, it estimates the reward probability of each option (**Fig. 1A**; Process 1). Then, based on this recognition, the agent selects an action based on reward and curiosity (**Fig. 1A**; Process 2).

In Process 1, we modeled the recognition process of reward probability using sequential Bayesian updating. The agent assumed that the reward is probabilistically generated from the latent cause *w* for both options (Fig. **1B**, left), and these probabilities are represented by a sigmoidal function of *w*_*i*_ as

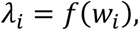

where *λ*_*i*_ and *w*_*i*_ denote the recognition of the reward probability of option *i* and its control variable, respectively, and *f*(*x*) = 1/(1 + *e*^−*x*^) (Fig. **1B**, middle). In other words, either reward or non-reward was observed at the probabilities *λ*_*i*_ and 1 − *λ*_*i*_ if the agent selected option *i*.

In addition, the agent assumed an environment in which the reward probabilities for both options could change ambiguously over time (**Fig. 1C, upper panel**). To describe this, *w* is fluctuated by a random walk as

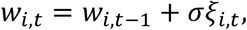

where *t, ξ*_*i,t*_, and *σ* denote the trial of the two-choice task, Gaussian noise with zero mean and unit variance, and the noise intensity, respectively. Accordingly, the reward probabilities for both options can be described by *λ*_*i,t*_ = *f*(*w*_*i,t*_). The temporal dynamics of the latent variable *w*_*i,t*_ and reward observation are described by the state-space model from the viewpoint of the agent (agent-SSM) (**Fig. 1C, middle**).

Given the above environment, the agent aims to estimate the reward probability *λ*_*i,t*_ of each option at each observation. To describe the confidence of *λ*_*i,t*_, the agent is assumed to estimate the control variable *w*_*i,t*_ as a distribution *P*(*w*_*i,t*_) and update its distribution. In general, *P*(*w*_*i,t*_) can be updated by action and reward observation *o*_*t*_ (**Fig. 1C, lower**), which can be roughly expressed as follows:

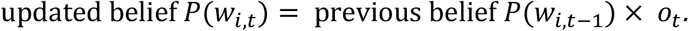

This process was formulated using Bayes’ rule (see Methods). We then derived the updates of the parameters as follows:

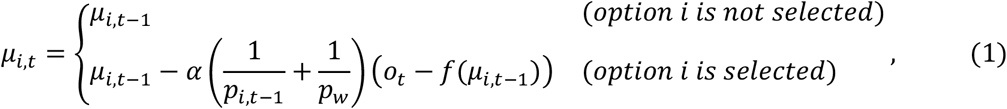

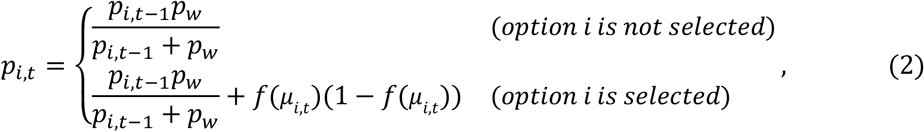

where *p*_*w*_ = 1/*σ*^2^; *μ*_*i,t*_ and *p*_*i,t*_ denote the mean and precision (i.e., inverse of variance) of *P*(*w*_*i,t*_) (see the details in Methods), respectively. If the *i*-th choice is not selected, its precision decreases (i.e., *p*_*i,t*+1_ < *p*_*i,t*_). With observation, belief is updated by the prediction error (i.e., *o*_*t*_ − *f*(*μ*_*i,t*_)), and its precision is improved. Because the reward probability is represented by *f*(*w*_*i,t*_), the estimated reward probability, that is, the recognized reward probability, should be transformed as *f*(*μ*_*i,t*_). Similarly, the confidence of the recognized reward probability should be evaluated not in *w*_*i,t*_ -space, but in 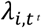-space, and hence the confidence is defined by *γ*_*t*_ = *p*_*i,t*_/*f*^′^(*μ*_*i,t*_)^2^ (see Methods).

In Process 2, the agent selects actions according to the recognition of the reward probability and its confidence in Process 1. Agents have two kinds of motivations in decision-making: the desire to maximize the reward, and curiosity—the desire to update their beliefs with more confidence (**Fig. 1D, upper**). This sum, called “expected utility” in this study, can be expressed as

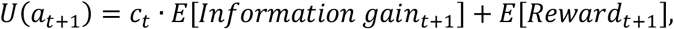

where “information gain” represents the information derived from a new observation based on previous recognition, *E*[*x*] denotes the expectation value of *x* based on current recognition, and *c*_*t*_ denotes a meta-parameter describing the intensity of curiosity, which weights the expected information gain. We regard *c*_*t*_ as varying over time (**Fig. 1D, lower panel**). We analytically derived *U*(*a*_*t*+1_) as a function of 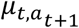 and 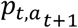 (see Methods). In decision-making, the agents selected action *a*_*t*+1_ with the highest expected utility with a higher probability as

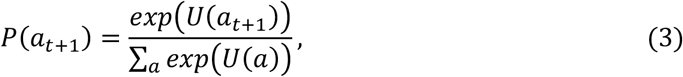

which is a SoftMax function.

### Sequential recognition and decision-making in the simulation

To validate our model, we performed simulations for two cases. In the first case, where reward probabilities were constant and different between the two options (**Fig. 2A**), the agent preferred to select the option with the higher reward probability (**Fig. 2B**). The recognized reward probabilities converged to ground truths, indicating that the agent accurately recognized the reward probabilities (**Fig. 2C**). The recognition confidence changed over time depending on the behavior; the confidence increased for the option that was selected more (**Fig. 2D**). The expected information gain was lower for the option with higher confidence (**Fig. 2E**), whereas the expected reward consistently followed the recognized reward probability (**Fig. 2F**). The expected utility, which is the sum of the expected information gain and reward, represents the value of each selection (**Fig. 2G**), resulting in the agent preferentially selecting the option with the higher expected utility.

**Fig. 2:**
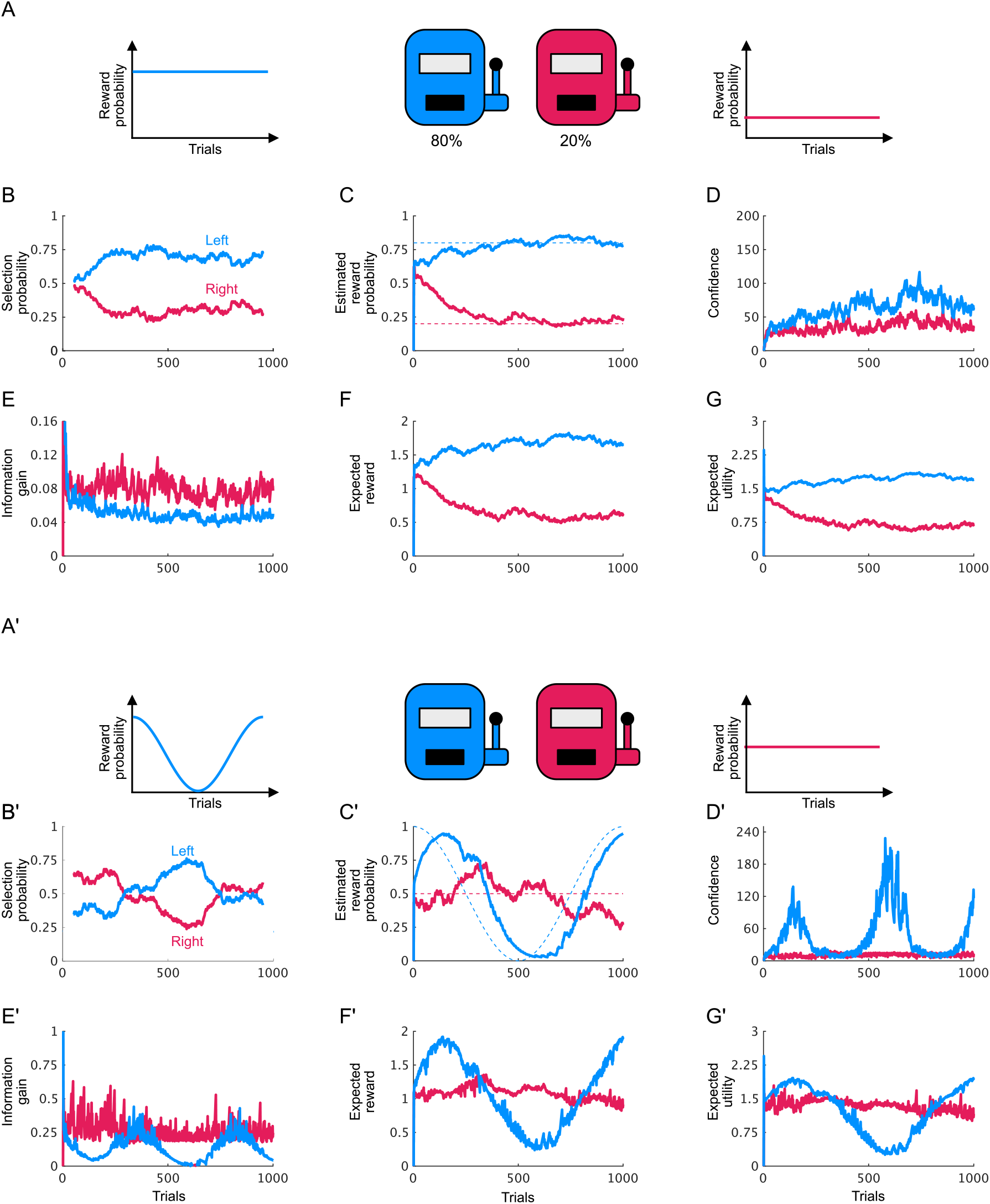
Simulations of the decision-making model. **(A)** Two-choice task with constant reward probabilities. **(B)** Selection probabilities for left and right options. These were plotted by moving average 101 window width. **(C)** Recognized reward probabilities for left and right options. Dashed lines represent the ground truths of the reward probabilities. **(D)** Confidences of reward probability recognitions for left and right options. Expected brief updates **(E)**, expected reward **(F)**, and expected utility **(G)** for left and right options. **(A’)** Two-choice task with constant and temporally varying reward probabilities for left and right options, respectively. **(B’-G’)** The same as (B-G). In these simulations, the curiosity parameter and reward seeking parameter were all kept constant at *c* = 1 and *P*_*o*_ = 0.9, respectively.

In the second case, we assumed a time-dependent reward probability (**Fig. 2A’**). In the simulation, the agent adaptively changed its recognition of the reward probability following the change in the true reward probability and selected the option with the higher estimated reward probability (**Fig. 2B’ and C’**). The confidence was also affected by the uncertainty of the reward probability at each time; the confidence was high where the reward probabilities were near-deterministic, around one and zero, and low where the reward probabilities were uncertain, approximately 0.5 (**Fig. 2D’**). The expected information gain of an option was negatively correlated with the confidence of the option (red line in **Fig. 2E’**), suggesting that the agent was curious about the uncertain option. As in the above case, the expected reward varied depending on the recognized reward probability (**Fig. 2F’**). The expected utility changed similarly to the expected reward, but the difference between the left and right options was less pronounced owing to curiosity (**Fig. 2G’**). These two demonstrations indicate that our model is able to represent the process of cognition and decision-making based on reward and curiosity.

### Curiosity-dependent irrational behaviors

Next, we examined how behavioral patterns are regulated by the intensity of curiosity and the degree of reward-seeking (**Fig. 3**). In a situation where the reward probabilities were zero for the left and 0.5 for the right (**Fig. 3A**), we simulated the model by varying the meta-parameters *c* and *P*_*o*_ (**Fig. 3B**). When the agent strongly desired the reward (*P*_*o*_ = 0.99, nearly equal to one) with no curiosity (*c* = 0), the agent preferred the right option with a higher reward probability (**Fig. 3C-(a)**). If the agent had no desire for a reward (*P*_*o*_ = 0.5) with high curiosity (*c* = 10), the agent preferred the right option with a higher reward probability (**Fig. 3C-(b)**). Although this behavior seems to be rational at first glance, the agent did not seek the reward, but rather sought the information, that is, belief update, driven by curiosity, resulting in preference for the uncertain option. When the agent moderately desired the reward (*P*_*o*_ = 0.5 to 1) with negative curiosity (*c* = −10), the agent conservatively selected the same option as the previous selection (**Fig. 3C-(c)**). As a result, the agent continuously selected either of the two options depending on the first selection. These results were expected.

**Fig. 3:**
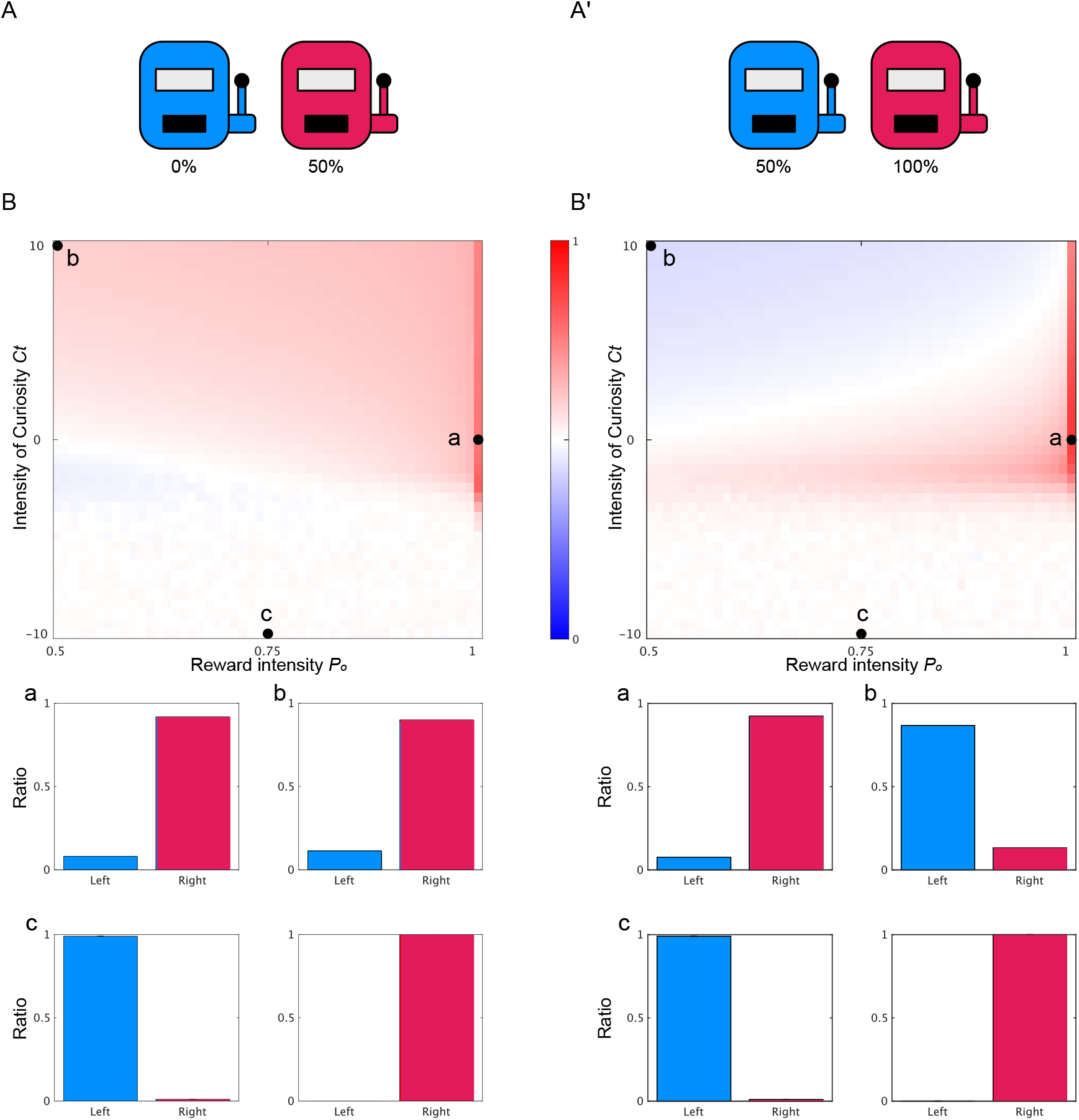
Curiosity-dependent irrational behaviors. **(A)** Two-choice task with different, constant reward probabilities: 0% for the left, and 50% for the right. **(B)** Heatmap of selection probability of the right option varying the parameters of curiosity and reward intensity (upper panel). There are three representative conditions indicated by black dots: *c* = 0, *P*_*o*_ = 1 [a], *c* = 10, *P*_*o*_ = 0.5 [b], and *c* = −10, *P*_*o*_ = 0.75 [c]. Lower four panels present the selection ratios of the right option for conditions [a], [b], and [c]. **(A’)** Two-choice task with different, constant reward probabilities: 50% for the left and 100% for the right. **(B’)** is the same as (B).

However, we obtained a non-trivial result in another situation, where the reward probabilities were 0.5 for the left and 1 for the right (**Fig. 3A’–C’**). As in the previous situation, the agents with a strong desire for the reward (*P*_*o*_ = 0.99, nearly equal to 1) preferred the right option with a higher reward probability (**Fig. 3C’-(a)**). The agent with no desire for reward (*P*_*o*_ = 0.5) and high curiosity (*c* = 10) preferred the left option with a lower reward probability (**Fig. 3C’-(b)**). This seemingly irrational behavior was the outcome of focusing on satisfying curiosity and not seeking rewards. In addition, as seen in the previous situation (**Fig. 3C-(c)**), the agents with negative curiosity (*c* = −10), irrespective of the desire for the reward, exhibited conservative selection (**Fig. 3C’-(c)**). Taken together, these results clearly indicate that behavioral patterns largely depend on the degree of conflict between reward and curiosity, which has been difficult to define because of ambiguity (**Fig. 3**).

### inverse FEP: Bayesian estimation of internal state

In the above cases, we assumed a constant balance between reward and curiosity. However, in reality, our feelings swing in a context-dependent manner. Although it is important to decipher the temporal swinging of the conflict between reward and curiosity in terms of neuroscience and psychology, it is difficult to quantify the conflict because of its temporal dynamics. Here, we developed a machine learning method called inverse FEP (iFEP) to quantitatively decipher the temporal dynamics of the internal state including curiosity meta-parameter from behavioral data.

In developing iFEP, we needed to switch the viewpoint from the agent to the observer; that is, from animals to us. In the agent-SSM, we described the sequential recognition of reward probabilities by the agent (**Fig. 1C, Fig. 4A,B**). Conversely, to determine the internal state of the agent, e.g., intensity of curiosity *c*_*t*_, recognition *λ*_*i,t*_ and its confidence *γ*_*i,t*_, we developed a state-space model from the observer’s eye (observer-SSM) (**Fig. 4B**). In the observer-SSM, the intensity of curiosity was assumed to change continuously over time, and the agent’s recognition of the reward probability was updated by equations (1) and (2) following FEP, but they were unknown to the observers. In addition, the agent’s actions were assumed to be generated depending on the intensity of its curiosity, recognition, and confidence, as described in equation (3), but the observers can only monitor the agent’s action and the presence of a reward. In iFEP, based on the observer-SSM, we estimate the latent internal state of agent *z* from observation *x* in a Bayesian manner as follows:

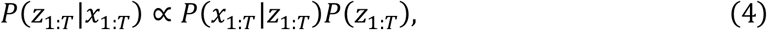

where *z*_*i*,1:*T*_ = {*μ*_*i*,1:*T*_, *p*_*i*,1:*T*_, *c*_1:*T*_}, *x*_1:*T*_ = {*a*_1:*T*_, *γ*_1:*T*_}, and the subscript 1: *T* indicates steps 1 to *T*. In this Bayesian estimation, a posterior distribution *P*(*z*_1:*T*_|*x*_1:*T*_) represents the observer’s recognition of the estimated *z*_1:*T*_ given observation *x*_1:*T*_ with uncertainty. A prior distribution *P*(*z*_1:*T*_) represents our belief, which is expressed as the ReCU model with random motion of the curiosity meta-parameter *c* as

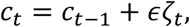

where *ζ*_*t*_ indicates white noise with zero mean and unit variance, and *ϵ* indicates its noise intensity. The likelihood *P*(*x*_1:*T*_|*z*_1:*T*_) represents the probability that *x*_1:*T*_ was observed assuming *z*_1:*T*_, which also follows the ReCU model. This Bayesian estimation, namely iFEP, was conducted using a particle filter and Kalman backward algorithm (see Methods).

**Fig. 4:**
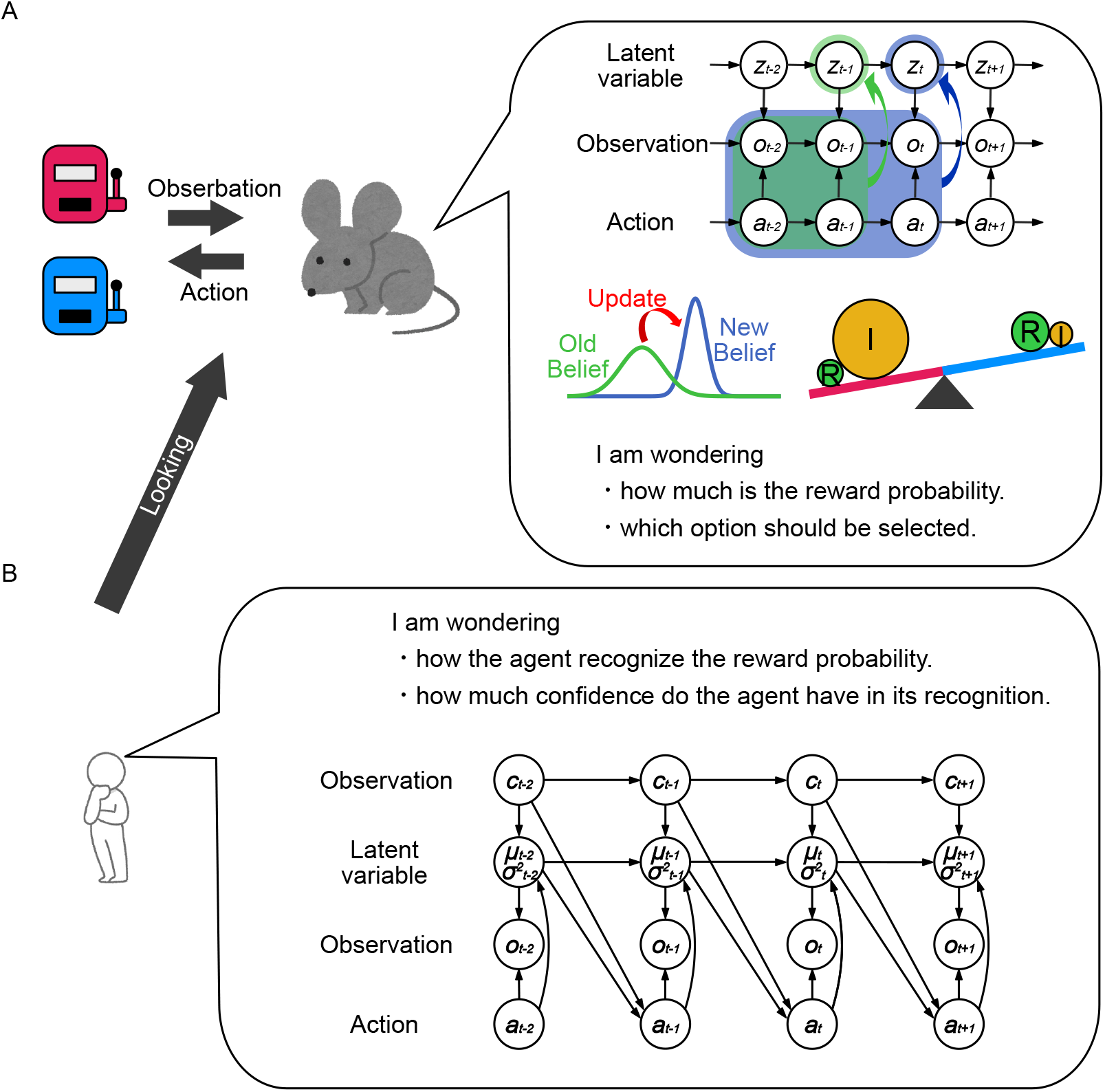
Scheme of inverse FEP by an observer of a decision-making agent. **(A)** An agent performing a two-choice task from the observer’s perspective. **(B)** Observer’s assumption about the agent’s decision-making. The agent is assumed to follow the decision-making model, as described in Fig. 1. **(C)** State-space model of the observer’s eye. For the observer, the agent’s reward– curiosity conflict, recognized reward probabilities, and their uncertainties are temporally varying latent variables, whereas the agent’s action and the presence/absence of a reward are observable. The observer estimates the latent internal states of the agent.

### Validation of iFEP

We tested the validity of the iFEP method by applying it to the artificial data generated by the ReCU model. Specifically, we simulated a model agent with non-constant curiosity in the two-choice task, where reward probabilities varied temporally. We then demonstrated that iFEP estimated the ground truth of the internal state of the simulated agent, that is, the agent’s intensity of curiosity, recognition, and confidence (**Fig. 5**). Therefore, iFEP must be powerful in clarifying decision-making processing and the accompanying temporal swing in the conflict between reward and curiosity.

**Fig. 5:**
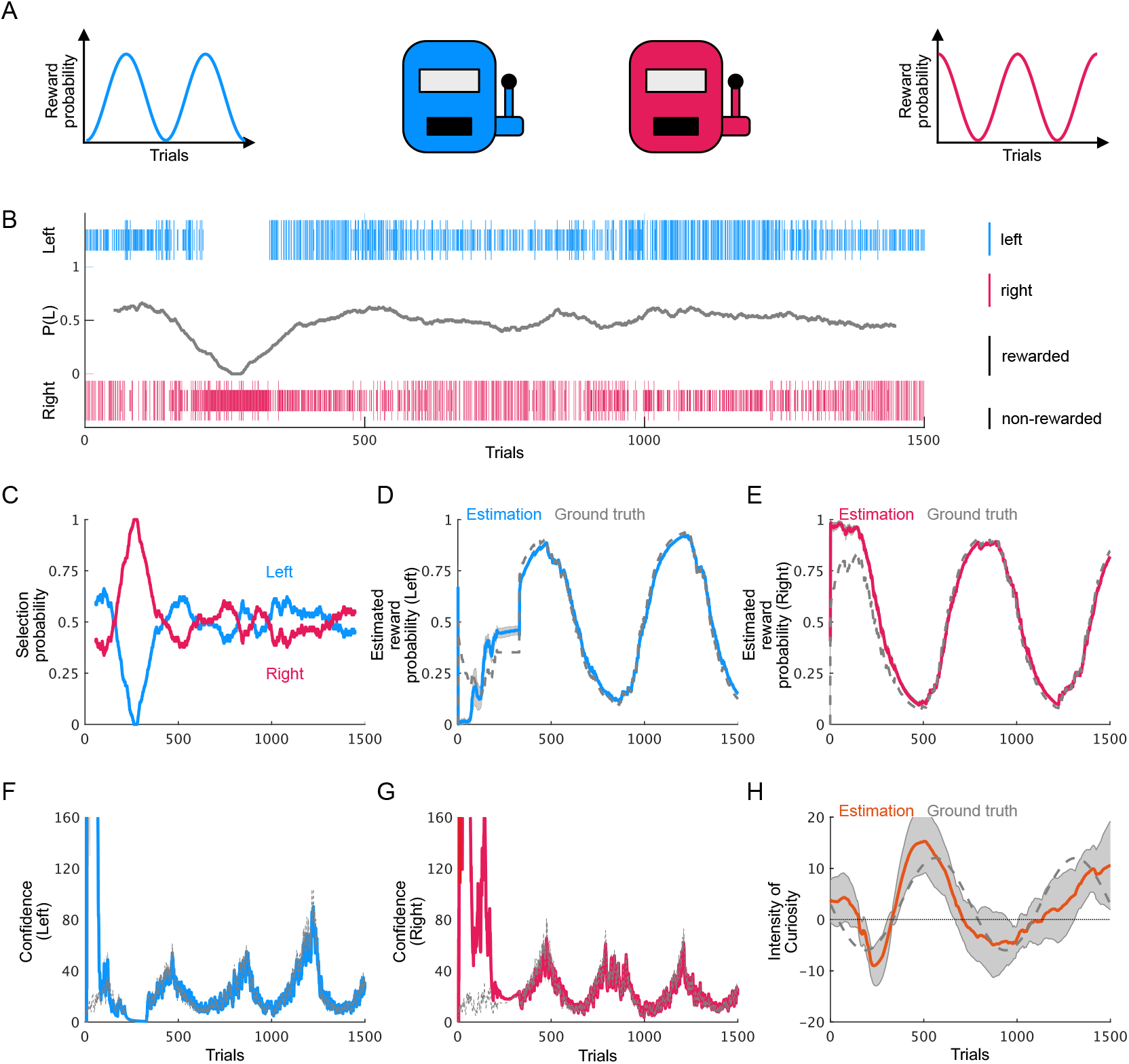
Estimation of the simulated agent’s internal state by inverse FEP. **(A)** Two-choice task with temporally varying reward probabilities and agent having temporally varying curiosity. **(B)** Simulated agent’s behaviors. Blue and red vertical lines indicate selections of left and right options, respectively. Long and short vertical lines indicate the presence and absence of rewards, respectively. Gray line indicates the moving average of selection probability of the left option with 101 window width. **(C)** Moving average of selection probabilities for left and right options. **(D– H)** Simulated agent’s behavior-driven estimations of agent-recognized reward probabilities for left (D) and right (E) options, agent’s confidence about recognized reward probabilities for left (F) and right (G) options, and agent’s curiosity (H). Orange and dashed blue lines represent the estimations and ground truths, respectively.

### Negative curiosity of rat behaviors decoded by iFEP

Finally, we applied iFEP to actual rat behavioral data from the two-choice task experiment with temporally varying reward probabilities (**Fig. 6A**)^17^. In this experiment, once the reward probabilities were suddenly changed in a discrete manner, the rat slowly adapted to select the option with the higher reward probability (**Fig. 6B**), suggesting that the rat sequentially updated its recognition of the reward probability. Based on these behavioral data of the rat, iFEP estimated the internal state, that is, the intensity of curiosity, the recognized reward probabilities, and their confidence levels (**Fig. 6C-E**). We found that the rat was not perfectly aware of the true reward probabilities but was able to recognize increases and decreases in reward probability (**Fig. 6C and D**). We also found that confidence increased with choice and decreased with no choice (**Fig. 6E and F**). Interestingly, the curiosity held by the rat was always estimated to be negative (**Fig. 6G**). In other words, the rat conservatively preferred certain choices but did not explore uncertain choices, regardless of the recognition of reward probability.

**Fig. 6:**
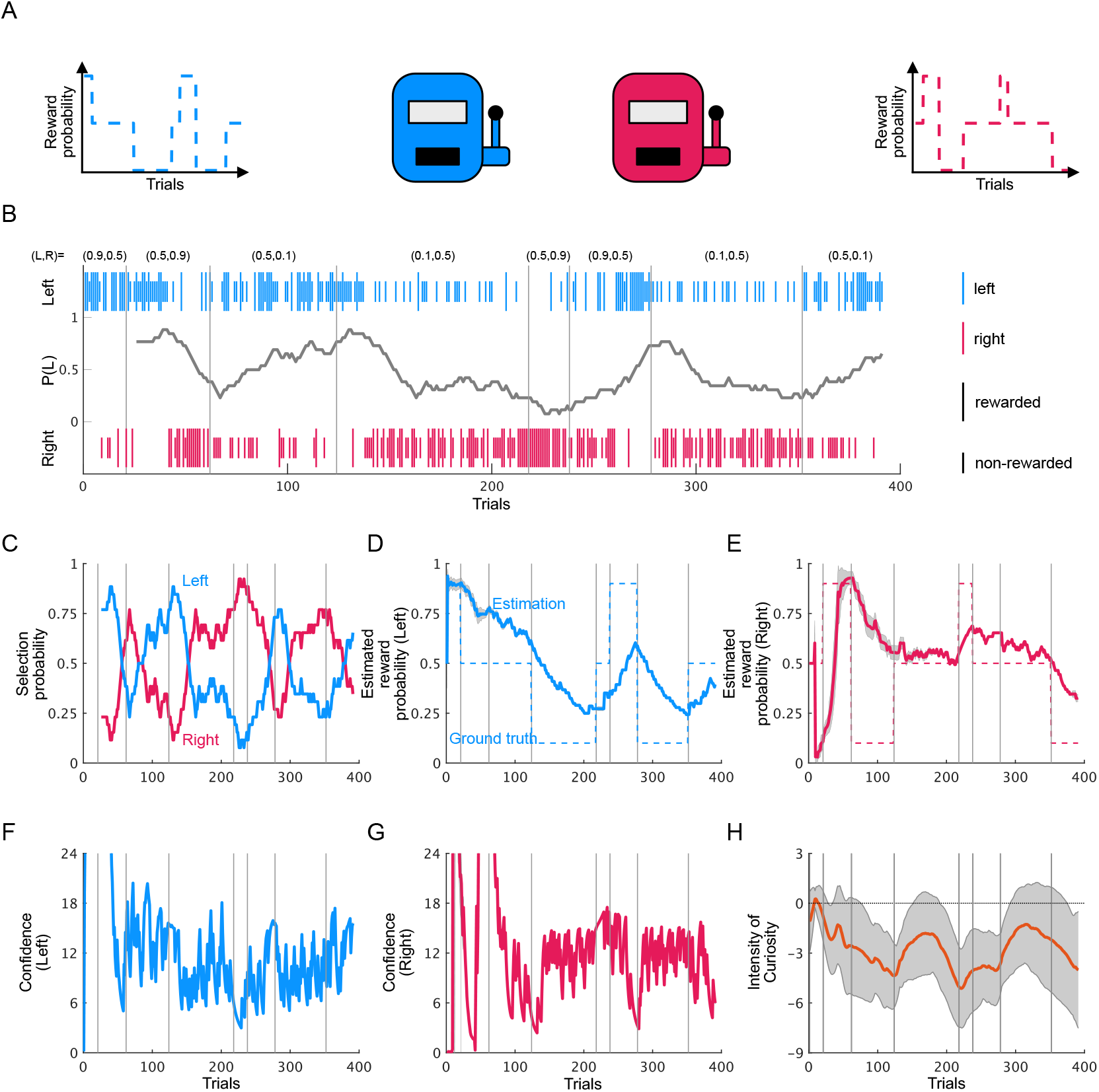
Estimation of the rat’s internal state by inverse FEP. **(A)** A rat in the two-choice task with temporally switching reward probabilities. **(B)** Rat’s behaviors in the two-choice task. A public dataset at https://groups.oist.jp/ja/ncu/data was used^5^. Blue and red vertical lines indicate selections of left and right options, respectively. Long and short vertical lines indicate the presence and absence of rewards, respectively. Gray line indicates the moving average of the selection probability of the left option with 25 window-width backward. **(C)** Moving average of selection probabilities for left and right options. **(D, E)** Rat behavior-driven estimations of agent-recognized reward probabilities for left (D) and right (E) options. Red and dashed blue lines represent the estimations and ground truths, respectively. **(F–H)** Rat behavior-driven estimations of agent’s confidence about recognized reward probabilities for left (F) and right (G) options, and agent’s curiosity (H).

To further show that negative curiosity was not estimated by chance, we conducted Monte Carlo testing (**Fig. S1**). As a null hypothesis, we considered an agent who has no curiosity and decides on a choice only depending on its recognition of the reward probability. Following the null hypothesis, we simulated a series of the agent’s choices using iFEP-estimated reward probability recognition and its confidence with zero curiosity (i.e., *c*_*t*_ = 0). The same simulations were repeated 1,000 times, and the curiosity was estimated using iFEP for each (**Fig. S1B**). We then plotted the null distribution for temporal average of estimated curiosity, as a test statistic (**Fig. S1B**). Compared to the null distribution, the temporal average curiosity estimated from the actual rat behavior was located to the left of the significance level (*p* = 0). These results suggest that rats have negative curiosity; they made choices based on confidence in reward probability recognition rather than the recognition itself.

## Discussion

Previous studies have discussed decision-making subject to rationality and optimality, in which irrational behaviors typical of animals cannot be tractable^18^. By contrast, we proposed the ReCU model, which expresses the mental conflict between reward and curiosity. In addition, we developed the iFEP method to decode the conflict between reward and curiosity based on the time-series behavioral data of a two-choice task.

What is the neural basis that controls psychological states, including mental conflict? This question cannot be addressed simply by comparing neural activities and animal actions because the psychological state is not reflected from each primitive action itself but must be behind a series of actions. Thus, it is important to define psychological states and estimate latent psychological states, such as curiosity, confidence, and reward prediction error, from animal behaviors, which enables us to compare the estimated psychological states and neural activities^12,15,19^. The iFEP method is an essential tool for this purpose. Therefore, the iFEP approach will contribute to the future development of neuroscience for psychological states.

Our approach has three novel characteristics compared with previous models. First, we addressed the irrationality of decision-making. The expected free energy was originally expressed as the sum of the expected information gain and the expected reward, both of which were equally weighted^10,11^, leading to Bayesian optimality. However, animals, including humans, perform decision-making under bounded rationality. Consequently, their computational performance is restricted by the limited size of the brain and time constraints for needing to make quick decisions. Thus, they do not necessarily achieve Bayesian optimization. By contrast, our study formulated irrational decision-making by introducing arbitrary weights on the expected information as a meta-parameter controlling curiosity. Therefore, our model represents individuality. Second, we addressed temporally changing curiosity, which might be situation-dependent. Animals and humans face a dilemma between reward and curiosity, which causes curiosity to vary over time. Third, we addressed an inverse problem using the iFEP method, which quantitatively decodes psychological states, including the intensity of curiosity, from the viewpoint of an observer of animals. This method is able to evaluate not only individual-specific properties, such as curious and conservative characters, but also temporal mental swings in a situation-dependent manner.

Decision-making on the two-choice task has been computationally addressed based on two types of modeling: RL and FEP. Studies based on RL described the temporal update of action values, which is called Q-learning^6^, but with a lack of recognition of the environment. In RL reinforcement modeling, it is common to represent the degree of exploration by a temperature parameter, which derives randomness in action selection but does not lead to information-seeking behavior, which corresponds to curiosity in our approach. Several studies have estimated the dynamics and parameters of Q-learning^15,17,19,20^. By contrast, Schwartenbeck et al. modeled decision-making behavior in a two-choice task based on FEP^21^. In their study, they succeeded in representing animal behaviors driven not only by reward but also by curiosity. However, they assumed that the intensity of curiosity was fixed at one (*c*_*t*_ = 1), at which the Bayes optimality was satisfied, and did not address its temporal variability as assumed and estimated in our iFEP approach. Furthermore, the related modeling differs from RL and FEP. Ortega and Braun formulated FEP, which describes irrational decision-making^22,23^. Interestingly, their formulation was based on microscopic thermodynamics, and the temperature parameters controlled this irrationality. However, the thermodynamics-based FEP did not treat the sequential update of the recognition from the observations.

Finally, it is worth discussing future perspectives of our iFEP approach in medicine. Generally, mental diagnosis relies on medical interviews and has not been quantitatively evaluated. Our iFEP method can quantitatively estimate the psychological state of patients based on their behavioral data. For example, patients with social withdrawal, also known as “Hikikomori,” have no interest in anything. In this case, social withdrawal would be characterized by a negative value of curiosity in our FEP model. Therefore, the iFEP method could be effective in diagnosing mental disorders.

## Methods

### FEP for reward probability recognition

In the reward probability recognition process, the agent assumes an environment expressed by a state-space model (**Fig. 1C**) as

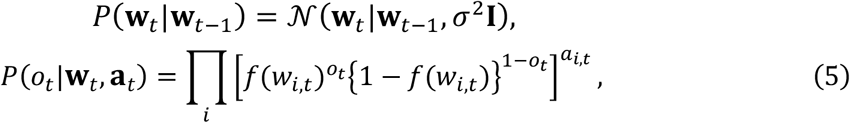

where **w**_*t*_ and *o*_*t*_ denote the latent variables controlling the reward probability of both options at step *t* (**w**_*t*_ = (*w*_1, *t*_, *w*_2, *t*_)^*T*^) and the observation of the presence of the reward (*o*_*t*_ ∈ {0,1}), respectively; **a**_*t*_ denotes the agent’s action at step *t*, which is represented by a one-hot vector (**a**_*t*_ ∈ {(1,0)^*T*^, (0,1)^*T*^}); 𝒩(**x**|**μ**, Σ) denotes the Gaussian distribution mean **μ** and variance Σ; *σ*^2^ denotes the variance of the transition probability of **w**; and 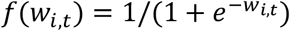, which represents the probability of reward of option *i* at step *t*. The initial distribution of **w**_1_ is given by *P*(**w**_1_) = *N*(**w**_1_|0, *k***I**), where *κ* denotes variance.

Here, we modeled the agent’s recognition process of reward probability using sequential Bayesian updating as follows:

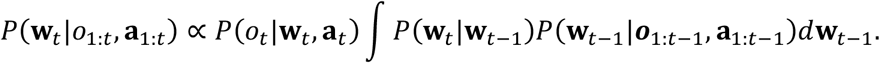

Because of the non-Gaussian *P*(*o*_*t*_|**w**_*t*_, **a**_*t*_), the posterior distribution of **w**_*t*_, *P*(**w**_*t*_|*o*_1:*t*_, **a**_1:*t*_), becomes non-Gaussian and cannot be calculated analytically. To avoid this problem, we introduced a simple posterior distribution approximated by a Gaussian distribution, as follows:

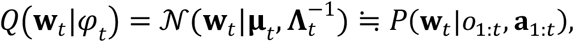

where *φ*_*t*_ = {**μ**_*t*_, **Λ**_t_}, and **μ**_*t*_ and **Λ**_t_ denote the mean and precision, respectively (**μ**_*t*_ = (*μ*_1,*t*_, *μ*_2,*t*_)^*T*^; **Λ**_t_ = *diag*(*p*_1,*t*_, *p*_2,*t*_)). *Q*(**w**_*t*_|*φ*_*t*_) denotes the recognition distribution. The model agent aims to update the recognition distribution through *φ*_*t*_ at each time step by minimizing the surprise, which is defined by−*lnP*(*o*_*t*_|*o*_1:*t*−1_). The surprise can be decomposed as follows:

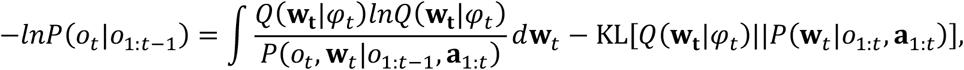

where KL[*q*(**x**)||*p*(**x**)] denotes the Kullback–Leibler (KL) divergence between the probability distributions *q*(**x**) and *p*(**x**). Because of the non-negativity of KL divergence, the first term is the upper bound of the surprise:

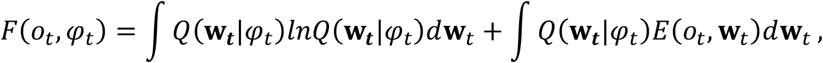

which is called the free energy, where *E*(*o*_*t*_, **w**_*t*_) = −*lnP*(*o*_*t*_, **w**_*t*_|*o*_1:*t*_, **a**_1:*t*_). The first term of the free energy corresponds to the negative entropy of a Gaussian distribution:

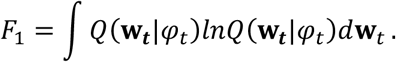

The second term is approximated as

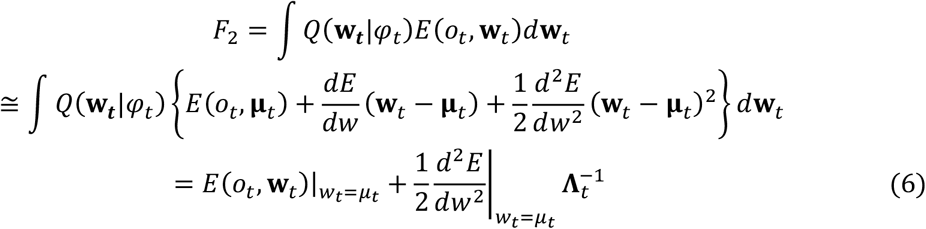

Note that *E*(*o*_*t*_, **w**_*t*_) is expanded by a second-order Taylor series around **μ**_*t*_. At each time step, the agent updates *φ*_*t*_ by minimizing *F*(*o*_*t*_, *φ*_*t*_).

### Calculation of free energy

The free energy is derived as follows. *F*_1_ simply becomes

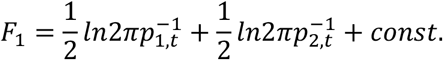

For computing *F*_2_,

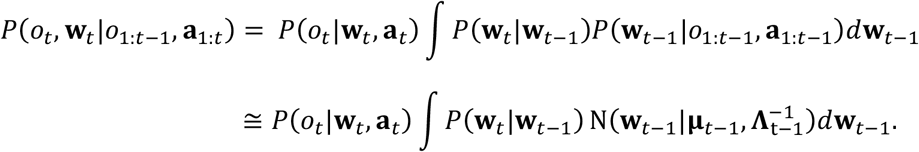

In the second line of this equation, we use the approximated recognition distribution as the previous posterior *P*(**w**_*t*−1_|*o*_1:*t*−1_, **a**_1:*t*−1_). This equation can be written as follows:

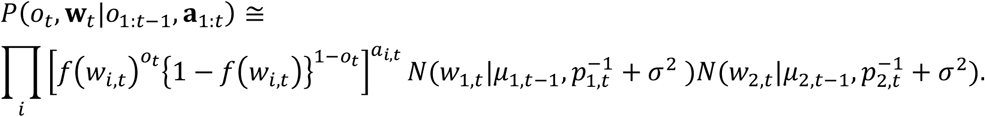

Then,

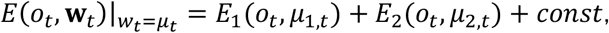

where

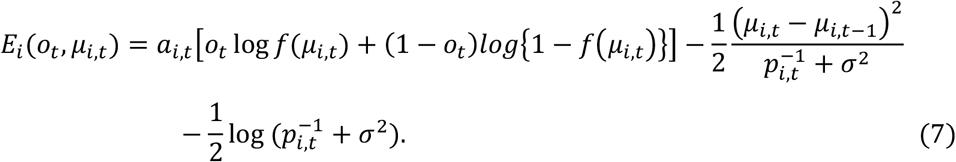

Thus, *F*_2_ is calculated by substituting equation (7) into equation (6). Taken together,

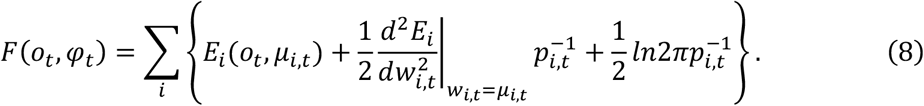

### Sequential updating of the agent’s recognition

The updating rule for *φ*_*t*_ was derived by minimizing the free energy. The optimized *p*_*i,t*_ can be computed by 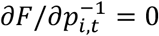, which leads to the following:

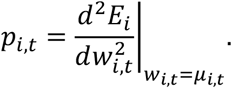

By substituting *p*_*i,t*_ into equation (8), the second term in the summation becomes constant, irrespective of *μ*_*i,t*_. Thus, *μ*_*i,t*_ is updated by minimizing only the first term as

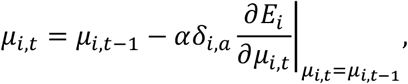

where *α* is the learning rate. These two equations finally lead to equations (1) and (2).

### Model for action selection

The agent selects a choice with higher expected utility defined by

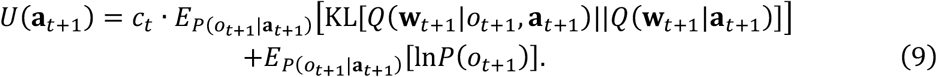

The first and second terms represent the expected information gain and expected reward, respectively, and *c*_*t*_ denotes the intensity of curiosity at time *t*.

In the first term, the posterior and prior distributions of **w**_*t*+1_ in the KL divergence are derived as

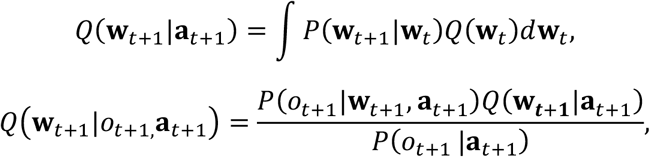

respectively. This KL divergence is called a Bayesian surprise, representing the extent to which the agents’ beliefs are updated by observation. Note that *o*_*t*+1_ is a future observation, therefore the first term was expected by *P*(*o*_*t*+1_ |**a**_*t*+1_), which can be calculated by

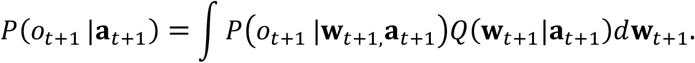

In the second term, the reward is quantitatively interpreted as the desired probability of *o*_*t*+1_. For the two-choice task, we use

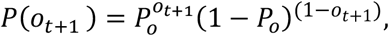

where *Po* indicates the desired probability of the presence of a reward. According to the probabilistic interpretation of reward^10,11^, the presence and absence of a reward can be evaluated by ln *P*_*o*_ and ln(1 − *P*_*o*_), respectively. Because rewards are relative, in this study, we set ln{*P*_*o*_/(1 − *P*_*o*_)} and 0 for the presence and absence of a reward, respectively.

### Calculation of the expected utility

Here, we present the calculation of the expected utility. The KL divergence in the first term of equation (9) can be transformed into

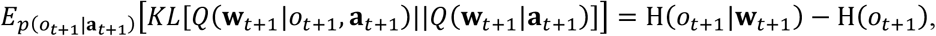

where the first and second terms represent the conditional and marginal entropies, respectively:

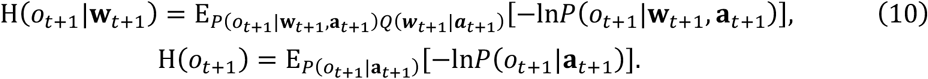

The conditional entropy H(*o*_*t*+1_|**w**_*t*+1_) can be calculated by substituting equation (5) into equation (10) as

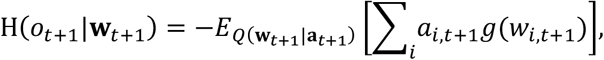

where

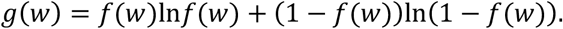

Here, we approximately calculate this equation by using the second-order Taylor expansion as

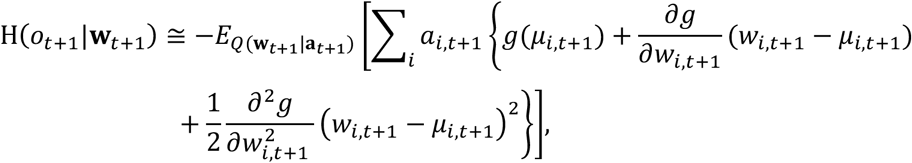

which leads to

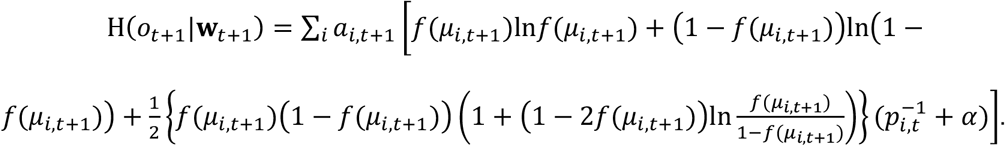

The marginal entropy H(*o*_*t*+1_) can be calculated as

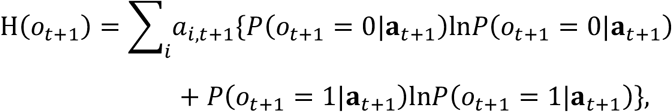

where

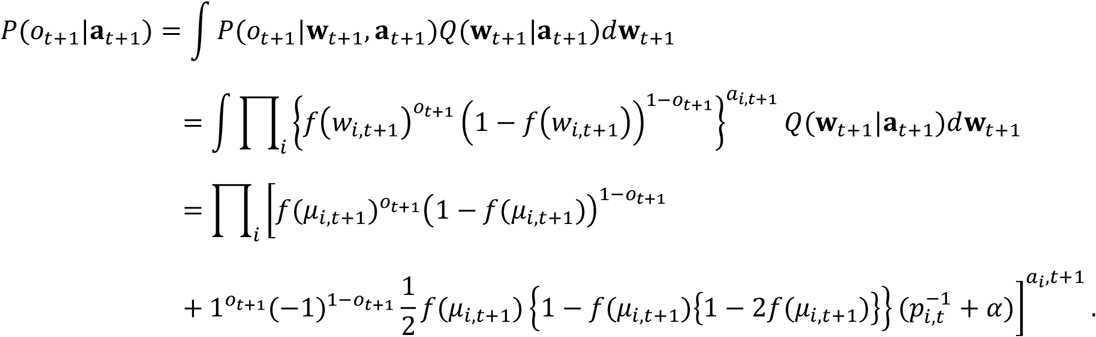

The second term of the expected utility (equation (10)) is calculated as

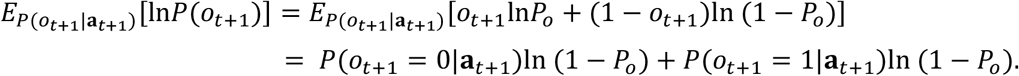

### The observer-SSM

We constructed the observer-SSM, which describes the temporal transitions of the latent internal state of agent *z* and the generation of action, from the viewpoint of the observer of the agent. This is depicted as a graphical representation in Fig. 4. As prior information, we assumed that the agent acts based on the internal state, that is, the intensity of curiosity, the recognized reward probabilities, and their confidence levels. The intensity of curiosity was assumed to change temporally as a random walk:

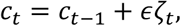

where *ζ*_*t*_ denotes white noise with zero mean and unit variance, and *ϵ* denotes its noise intensity. Other internal states, that is, *μ*_*i*_ and *p*_*i*_, were assumed to update as equations (1) and (2). The transition of the internal state is expressed by the probability distribution,

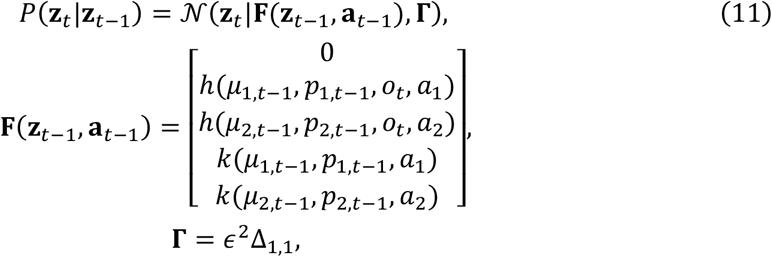

where 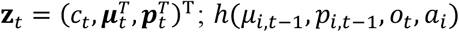 and *k*(*μ*_*i,t*−1_,*p*_*i,t*−1_,*o*_*t*_,*a*_*i*_) and *k*(*μ*_*i,t*−1_,*p*_*i,t*−1_,*a*_*i*_) represent the right-hand sides of equations (1) and (2), respectively, and Δ_*i,j*_ denotes a variance-covariance matrix with 1 at (*i, j*) component and 0 at others. In addition, the agent was assumed to select an action **a**_*t*+1_ based on the expected utilities, as follows:

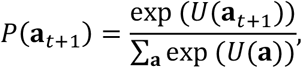

and the reward was obtained by the following probability distribution:

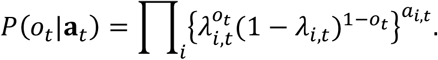

### iFEP by particle filter and Kalman backward algorithm

Based on the observer-SSM, we estimated the posterior distribution of the latent internal state of agent *z*_*T*_ given all observations from 1 to *T* (*x* _1:*T*_) in a Bayesian manner, i.e., *P*(*z*_*t*_|*x* _1:*T*_). This estimation was done by forward and backward algorithms, which are called filtering and smoothing, respectively.

In filtering, the posterior distribution of *z*_*t*_ given observations until *t* (*x* _1:*T*_) is sequentially updated in a forward direction as

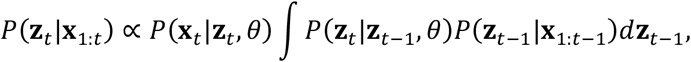

where 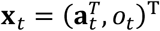 and *θ* = {*σ*^2^, *α, P*_*o*_}. The prior distribution of **z**_1_ is

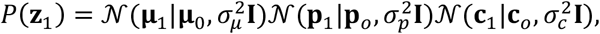

where **μ**_0_, **p**_*o*_and **c**_0_ denote means, and 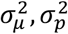 and 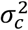 denote variances. We used a particle filter^24^ to sequentially calculate the posterior *P*(**z**_*t*_|**x** _1:*t*_), which cannot be analytically derived because of the nonlinear transition probability.

After the particle filter, the posterior distribution of **z**_*t*_ given all observations (**x** _1:*T*_) is sequentially updated in a backward direction as

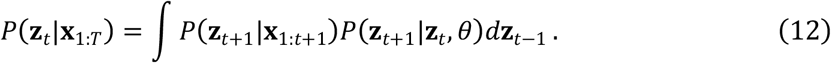

However, this integration is intractable due to non-Gaussian *P*(**z** _*t*−1_|**x**_1:*t*−1_), which was represented by particle ensemble in the particle filter, and non-linear relationship between **z**_*t*−1_ and **z**_*t*_ in *P*(**z**_*t*_|**z**_*t*−1_, *θ*) (equation (11)). Thus, we approximated *P*(**z**_*t*_|**x**_*t*:1_) as 𝒩(**z**_*t*_|**m**_*t*_, **V**_*t*_), where **m**_*t*_ and **V**_*t*_ denote sample mean and sample variance of particles at *t*, whereas we linearized *P*(**z**_*t*_|**z**_*t*−1_, *θ*) as

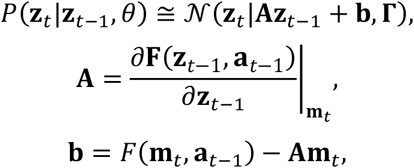

where **A** denotes Jacobian matrix. Because these approximations make integration of equation (12) tractable, the posterior distribution *P*(**z**_*t*_|**x**_1:*T*_) can computed by Gaussian distribution as

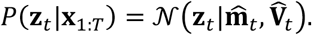

Its mean and variance was analytically updated by Kalman backward algorithms^25^:

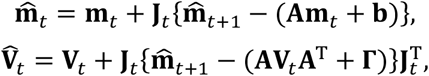

where

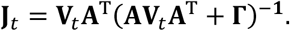

### Estimation of parameter in iFEP

To estimate parameter *θ*, we extended the observer-SSM to a self-organizing state-space model^26^ in which *θ* was also addressed as constant latent variables:

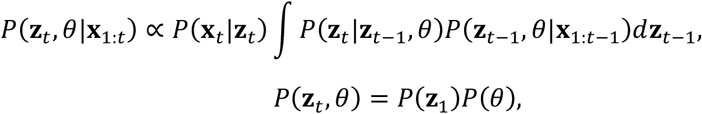

where 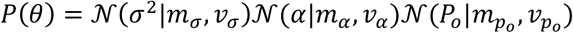. To sequentially calculate the posterior *P*(**z**_*t*_, *θ*|**x**_1:*t*_) using the particle filter, we augmented the state vector of all particles by adding parameter *θ*, which was not updated from randomly sampled initial values.

### Statistical testing with Monte Carlo simulations

In Fig. S1, we statistically tested negative curiosity estimated in Fig. 6. A null hypothesis is that an agent who has no curiosity (i.e., *c*_*t*_ = 0) and decides on a choice only depending on its recognition of the reward probability. Under the null hypothesis, model simulations were repeated 1,000 times under the same experimental conditions as in Fig. 6 and the curiosity was estimated for each using iFEP. We adopted temporal average of estimated curiosity as a test statistic and plotted the null distribution of the test statistic. Compared with the estimated curiosity of the rat behavior, we then computed p-value for one sided left tailed test.

## Acknowledgments

We are grateful to Prof. Kenji Doya and Dr. Makoto Ito for providing rat behavioral data. We thank organizers of the tutorial on the free-energy principle in 2019, that inspired this research. This study was supported in part by Grant-in-Aid for Transformative Research Areas (B) [grant number 21H05170], AMED [Grant Number JP21wm0425010], Moonshot R&D–MILLENNIA Program [grant number JPMJMS2024-9] by JST, the Cooperative Study Program of Exploratory Research Center on Life and Living Systems (ExCELLS) [program number 21-102], and the grant of Joint Research by the National Institutes of Natural Sciences [NINS Programme No. 01112102].

## Author contributions

H.N. conceived of the project. Y. K. and H.N. developed the method and Y.K. implemented the model simulation. Y.K. and H.N. wrote the manuscript.

## Data availability

We used the rat behavioral data published by Ito and Doya, *J Neurosci* (2009)^17^, which is publicly available at Prof. Doya Kenji’s homepage: https://groups.oist.jp/ja/ncu/data. Source codes will be open after publication.

## Supplementary Information

**Fig. S1:**
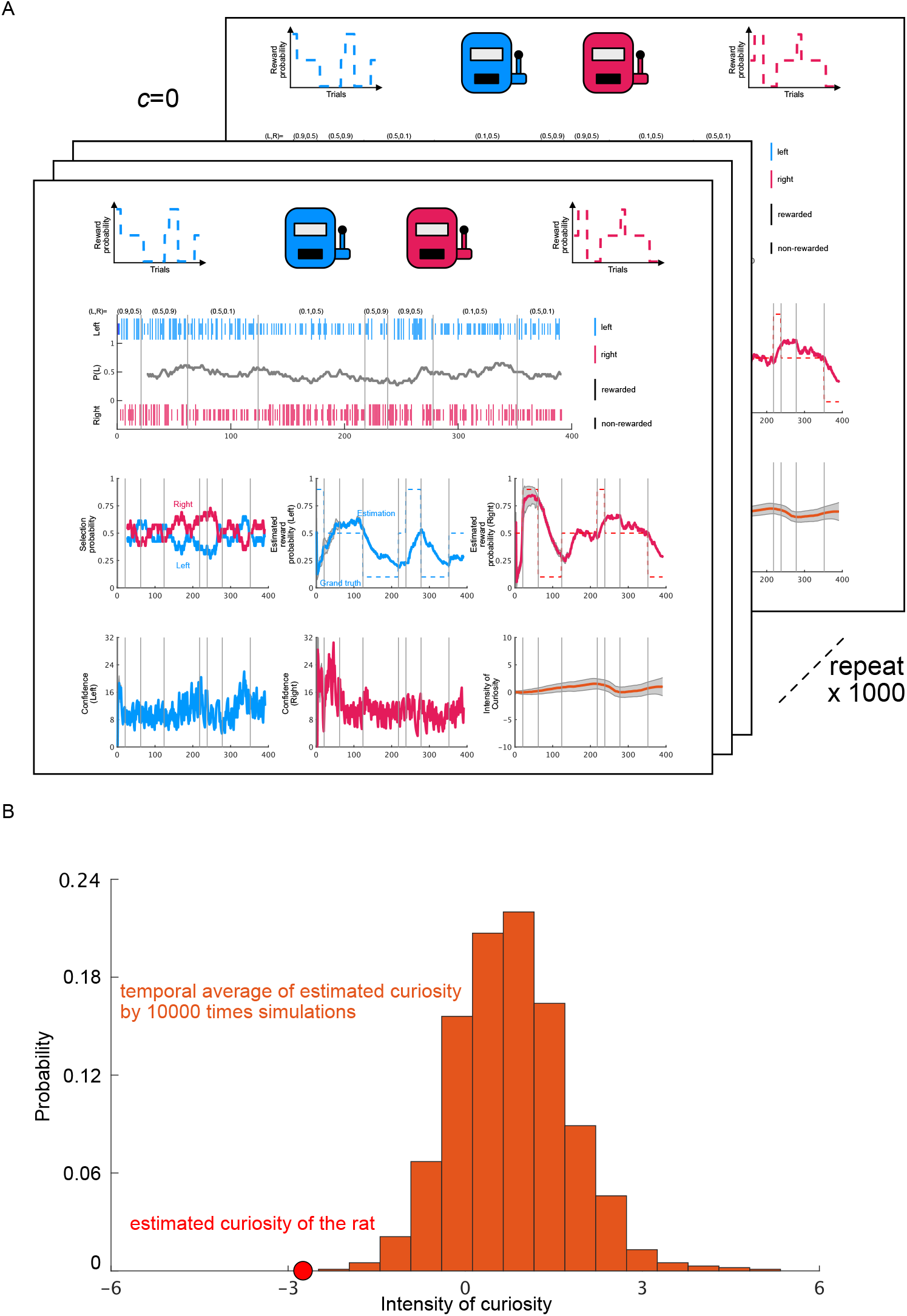
Monte Carlo statistical testing of estimated negative curiosity in rat. **(A)** Model simulations conducted under the same experimental conditions as in Fig. 6 under a null hypothesis that the rat behavior is generated by zero curiosity. Simulations were repeated 1,000 times, and the curiosity was estimated using iFEP for each. **(B)** Null distribution for temporal average of estimated curiosity by 1,000 times simulations. The estimated curiosity of the rat shown in red circles was on the left side and did not overlap with the distribution, which means *p*=0 (one-sided, left tailed test).

